# Mobile element-mediated carbapenem resistance in *Enterobacter hormaechei* in a Nigerian intensive care unit

**DOI:** 10.64898/2026.04.09.712135

**Authors:** Ifeanyi E. Mba, Erkison Ewomazino Odih, Olukemi Adekanmbi, Anderson O. Oaikhena, Gabriel T. Sunmonu, Ini Adebiyi, Adebimpe T. Gbaja, Olabisi G. Animashaun, Precious E. Osadebamwen, Olusola Idowu, David M Aanensen, Iruka N Okeke

## Abstract

Carbapenem-resistant Gram-negative bacteria pose a critical public health threat. The role of mobile genetic elements in driving their transmission and persistence remains poorly defined. In 2022, we investigated a suspected outbreak of carbapenem-resistant *Acinetobacter baumannii* (CRAB) in a Nigerian adult intensive care unit (ICU), using short-read whole genome sequencing (WGS) of carbapenem-resistant clinical and environmental isolates during the cluster period. Mobile element dynamics were then inferred from hybrid assemblies of Illumina and Oxford Nanopore reads. The suspected CRAB outbreak was ruled out by WGS but a carbapenem-resistant *Enterobacter hormaechei* ST114 bloodstream isolate was found to be indistinguishable from two environmental isolates, all recovered during the *Acinetobacter* surge. Hybrid assemblies revealed a strikingly conserved ∼19 Kb resistance island shared across all ST114 genomes. The island contained a *bla*_NDM-5_ cassette alongside many other antimicrobial resistance genes, within class 1 integronns and flanked by insertions sequences, located on a 46,176 bp plasmid. Using the ST114 plasmid’s hybrid assembly as scaffold, the same plasmid was identified in the genome of a *Klebsiella pneumoniae* ST15 isolate from the ICU environment during the same period. Additionally, re-interrogation of genomic surveillance data uncovered four clonal 2020 ST109 *Enterobacter* bloodstream isolates from the same facility that carried the resistance genes in the same context on a large 267,242 bp plasmid. Carbapenem resistance in hospital Enterobacterales is driven by both clonal expansion and horizontal spread of mobile resistance elements. These findings underscore the need to track mobile elements alongside bacterial lineages to inform evidence-based infection control, especially in low-resource settings.

**Impact Statement:** Carbapenem resistance among Enterobacterales remains a major public health threat, yet how mobile genetic elements contribute to their persistence and spread in hospital settings is still poorly understood. In this study, we investigated a suspected outbreak of carbapenem-resistant *Acinetobacter baumannii* in an adult intensive care unit in Nigeria. Although the outbreak was eventually ruled out, genomic analysis has shown the importance of careful interpretation of suspected outbreak cases in hospital settings. Our findings highlight the importance of close monitoring of ICU environments, the implementation of blood culture–based diagnostics, and the value of genomic support in outbreak investigations. These findings demonstrate that carbapenem resistance in hospital Enterobacterales is driven not only by clonal expansion but also by the horizontal dissemination of a highly stable *bla*_NDM-5_-associated MDR island capable of integrating into diverse plasmid backbones. This study emphasizes the need for genomic surveillance that tracks both mobile elements and bacterial lineages to strengthen outbreak investigations, especially in low-resource settings. It further underscores the links between clinical and environmental AMR reservoirs and reinforces the value of a One Health approach to controlling carbapenem resistance.

**Data summary:** FASTQ sequences were deposited in the NCBI BioSample database under accession numbers SAMN55915584 – SAMN55915597.

## Introduction

Carbapenems are last-resort antibiotics for treating multidrug-resistant Gram-negative bacterial infections (1). Carbapenem-resistant Gram-negative bacteria are of growing concern worldwide, particularly in healthcare settings and give rise to infections with high mortality rates (2). There remains a huge gap in our understanding of their epidemiology in Africa, where there are limited support and resources for genomic surveillance. Health care associated infections are frequent in Africa but less commonly detected, which has important implications for patient safety, although substantial heterogeneity limits direct comparisons (3).

In Europe, carbapenem-resistant Enterobacterales (CRE), including *Klebsiella pneumoniae, Escherichia coli*, and *Enterobacter* pose a significant threat to patients and healthcare systems (4), as does carbapenem-resistant *Acinetobacter baumanii* (CRAB). The impact of CRAB and CRE are more pronounced in low- and middle-income countries (LMICs), where carbapenems are out of reach for many patients and alternatives are often not available (5). Both CRAB and CRE can transfer genetic material to other pathogens within a given niche through horizontal gene transfer (HGT) [6] with possible implications in genome reorganization and long-term evolutionary consequences.

University College Hospital was a pilot site for *A Clinically Oriented Antimicrobial Resistance Network* (ACORN2) (6). As part of ACORN’s case-base surveillance we performed bimonthly HAI scanning of wards included in the pilot (7). We noticed an upsurge in HAIs in the adult ICU, co-incident with recovery of *Acinetobacter* isolates. This prompted whole genome sequencing of patient and environmental isolates recovered during the period. The investigation ruled out the suspected *Acinetobacter* outbreak but identified an *Enterobacter hormaechei* clone with a carbapenem resistance mobile element. Here we present findings of the genomic analysis and evidence pointing to a temporal and taxonomic broad range for the mobile element.

## Materials and Methods

### Strains

Blood samples from adult ICU patients were aseptically collected in BACTEC blood culture bottles and cultured using the automated BACTEC FX 40 machine. Broth from positive vials was Gram-stained and sub-cultured on blood and chocolate agar. Environmental samples were collected using sterile swabs and exposure agar (opened) plates. The swab samples were then cultured on appropriate culture media. All the cultured plates were incubated at 37 °C for 18-24 hours. Identification of isolates was performed using the Analytic Profile Index 20E (API-20E). Antibiotic susceptibility was determined using the Kirby-Bauer disc diffusion method, and carbapenem resistance was identified by resistance to meropenem (30µg) antibiotic disks based on the current CLSI guideline. Clinical and environmental isolates from the UCH ICU were sent to the national reference laboratory by the institution’s Infection Prevention and Control Committee. All isolates were re-identified on receipt, employing the VITEK2 system (GN-2413074203 cards), according to manufacturer’s recommendations and maintained at -80°C in LB:glycerol 1:1. *Escherichia coli* ATCC 25922 and *Acinetobacter baumanii* NTCC7363 were used as controls for media preparation and microbiology characterization.

### Whole-Genome Sequencing

Genomic DNA was extracted from the isolates using the commercially available Wizard DNA kits, (Promega; Wisconsin, USA), following the manufacturer’s instructions. Extracted DNA was quantified using a Qubit fluorometer with the dsDNA Broad Range assay (Invitrogen, California, USA). DNA libraries were prepared using the NEBNext Ultra II FS DNA Library Prep Kit for Illumina with 96 unique indexes (New England Biolabs, Massachusetts, USA; Cat. No. E6609L). Library concentrations were measured using the dsDNA High Sensitivity assay on a Qubit fluorometer (Invitrogen, California, USA), and average fragment sizes were assessed using a 2100 Bioanalyzer (Agilent). Libraries were sequenced on an Illumina MiSeq platform (Illumina, California, USA). Whole-genome sequencing data were generated for all isolates using illumina platform.

Nanopore sequencing was also performed on the *Enterobacter* strains using the Oxford Nanopore technology to generate complete genome assemblies for comprehensive analyses. High-molecular-weight genomic DNA was extracted using the FastDNA Spin Kit for Soil (MP Biomedicals, Santa Ana, CA, USA) according to the manufacturer’s protocol to minimise DNA fragmentation. Long-read sequencing libraries were prepared using the Ligation Sequencing Kit (SQK-LSK109) and sequenced on a MinION flow cell (R9.4.1) using MinKNOW v22.08.9 (Oxford Nanopore Technologies, Oxford, United Kingdom). Super-accuracy basecalling and demultiplexing of the raw reads were performed using MinKNOW v22.08.9.

### Bioinformatic analyses

Raw Illumina sequence reads were first processed to remove adapters and low-quality bases using Trimmomatic v0.38 (8). Trimmed reads were assembled de novo with SPAdes v3.12.0 (9). The quality of the raw sequence reads was assessed using FastQC v0.11.6 (10) (Supplementary Table 1). Genome assemblies were evaluated using QUAST v5.2.0 (https://github.com/ablab/quast). Species identification was performed using Bactinspector v0.1.3 (https://gitlab.com/antunderwood/bactinspector). Assembled genomes were screened for contamination using ConFindr v0.8.2 (11). Multilocus sequence typing (MLST) was performed using Pathogenwatch (12).

**Table 1:** Characteristics of clinical and environmental bacterial isolates sequenced during the HAI surge and their genomes.

| Isolates ID | Source | VITEK2 ID | WGS species designation | ST | Genome length | No. of contigs | N50 | G+C content | Resistance genes | Plasmid replicons |
| --- | --- | --- | --- | --- | --- | --- | --- | --- | --- | --- |
| UCH_0887_ABA | Patient | *Acinetobacter baumannii* | *Acinetobacter baumannii* | 2833 | 4,011,447 | 151 | 93,669 | 39.1 | 8 | 0 |
| UCH_B0199 | Patient | *Acinetobacter baumannii* | *Acinetobacter baumannii* | 919 | 4,187,791 | 92 | 151,998 | 39.0 | 14 | 0 |
| UCH_B0249 | Patient | *Acinetobacter baumannii* | *Proteus mirabilis* | - | 3,975,766 | 33 | 267,276 | 38.7 | 10 | 1 |
| UCH_778 | Patient | *E. cloacae* complex | *Enterobacter hormaechei* | 114 | 5,038,308 | 189 | 263,501 | 54.9 | 24 | 5 |
| UCH_710 | Environmental | *E. cloacae* complex | *Enterobacter hormaechei* | 114 | 5,057,769 | 140 | 507,693 | 54.9 | 24 | 5 |
| UCH_770 | Environmental | *E. cloacae* complex | *Enterobacter hormaechei* | 114 | 5,080,529 | 94 | 1,039,168 | 54.8 | 24 | 5 |
| UCH_619 | Environmental | *K. pneum pneumoniae* | *Klebsiella pneumoniae* | 3133 | 5,258,691 | 89 | 126,723 | 57.4 | 6 | 2 |
| UCH_935D_S13 | Environmental | *K. pneum pneumoniae* | *Klebsiella pneumoniae* | 395 | 5,549,563 | 85 | 276,030 | 57.2 | 15 | 6 |
| UCH_943_ECO | Environmental | *K. pneum pneumoniae* | *Klebsiella pneumoniae* | 967 | 5,572,066 | 68 | 377,798 | 57.0 | 13 | 5 |
| UCH_943_LLFB | Environmental | *K. pneum pneumoniae* | *Klebsiella pneumoniae* | 967 | 5,499,598 | 252 | 61,883 | 57.0 | 11 | 4 |
| UCH_B0251_KLEB | Environmental | *K. pneum pneumoniae* | *Klebsiella pneumoniae* | 14 | 5,594,104 | 136 | 296,409 | 57.1 | 14 | 3 |
| UCH_B0265_KLEB | Environmental | *K. pneum pneumoniae* | *Klebsiella pneumoniae* | 147 | 5,662,862 | 73 | 348,574 | 57.0 | 14 | 5 |
| UCH_P0485_KLEB | Environmental | *K. pneum pneumoniae* | *Klebsiella pneumoniae* | 15 | 5,863,190 | 162 | 157,670 | 56.7 | 22 | 5 |
| UCH_P0405_S11 | Environmental | *K. pneum pneumoniae* | *Escherichia coli* | 10 | 4,789,984 | 89 | 223,095 | 50.8 | 11 | 4 |

Oxford Nanopore long reads from select *Enterobacter* genomes were trimmed to remove adapters and low-quality regions using Porechop v0.2.4 (https://github.com/rrwick/Porechop). Long-read-only assemblies were generated using Flye v2.9 (13). Hybrid genome assemblies were generated using Unicycler v0.5.1 (14). Quality metrics of the short-read, long-read, and hybrid assemblies are shown in Supplementary Table 1. The assembled genomes were annotated using Bakta (15). Phylogenomic analysis was performed using GTDB-Tk v2.4.1 with the classify workflow [16].

Plasmid replicons were identified using PlasmidFinder v3.0.1 (16). Complete plasmid sequences were identified and extracted from the *E. hormaechei* genomes with complete assemblies (obtained via hybrid assembly) and used as scaffolds for the reconstruction of corresponding plasmids in the *K. pneumoniae* genomes. AMR genes were identified using ABRicate v1.0.1 (https://github.com/tseemann/abricate), which performs BLASTn searches against curated AMR gene databases. The NCBI AMRFinder database (17) was used, applying minimum identity and coverage thresholds of 90% and 80%, respectively. The plasmids from the *Enterobacter* hybrid assembly were typed using MOB-suite v3.1.9 [18]. Mobile genetic elements were identified using MobileElementFinder (v2.0.3) and VRprofile2 (https://tool2-mml.sjtu.edu.cn/VRprofile) with default parameters. MobileElementFinder enabled precise detection of individual insertion sequences, whereas VRprofile2 characterized broader and complex mobile regions, including mosaic resistance regions and AMR-associated mobile element clusters.

Single nucleotide polymorphisms (SNPs) were identified using Snippy v4.6.0 (18). Illumina paired-end reads from the two *Acinetobacter* isolates were mapped to the complete reference genome of *A. baumannii* ATCC 17978, while UCH770 was used as the reference for the *Enterobacter* strains. The reads from each isolate were mapped to the respective reference genome using Snippy’s default mapping and variant calling pipeline, which employs BWA-MEM v0.7.17 for alignment and FreeBayes v1.3.5 for SNP calling. Snippy output variant call files (VCF) were filtered using the default minimum coverage (10X) and minimum base quality (Q20) thresholds. Pairwise SNP distances between strains were calculated using snp-dists v0.8.2 [19] https://github.com/tseemann/snp-dists. To visualize gene organization and conserved synteny, the multidrug-resistance (MDR) regions harboring mobile genetic elements from the *Enterobacter* ST114 strains (UCH_770, UCH_778 and UCH_710) were extracted from their assembled genomes in GenBank format and compared. Gene cluster alignments and synteny plots were generated using Clinker v0.0.23 and Clustermap.js v0.1.0 (19) with default parameters, enabling visualization of conserved and variable gene arrangements across the MDR regions. Resulting figures were subsequently refined and annotated using Inkscape v1.4.3.

## Results

### Bacterial species recovered from the ICU

A consultant infectious disease specialist reported an unusually high number of suspected bacteriaemia infections in the Adult Intensive Care Unit (ICU) in May 2022, which was also reflected as a pronounced spike in HAI prevalence on the ACORN2 dashboard (Figure 1). Following a baseline prevalence of approximately 2–3%, the rate surged sharply, exceeded the 5% threshold and peaked at around 25%. By the end of the observation period, HAI prevalence had decreased to approximately 12%, but still remained above the baseline threshold indicated by the red reference line (Figure 1). Even though, due to resource limitations, only a fraction of patients that require blood cultures receive them, the laboratory reported that *Acinetobacter*, typically uncommon, had been cultured from four patients during the uptick, and were carbapenem-resistant, leading to suspicion of a CRAB outbreak. VITEK2 re-identification and Illumina whole genome sequencing of the presumptive *Acinetobacter* isolates and resistant isolates collected during environmental surveillance revealed that while two of the four presumptive *Acinetobacter* isolates were *Acinetobacter baumanii*, the other two were mis-identified *Proteus mirabilis* and *Enterobacter hormaechei* strains (Table 1).

**Figure 1.**
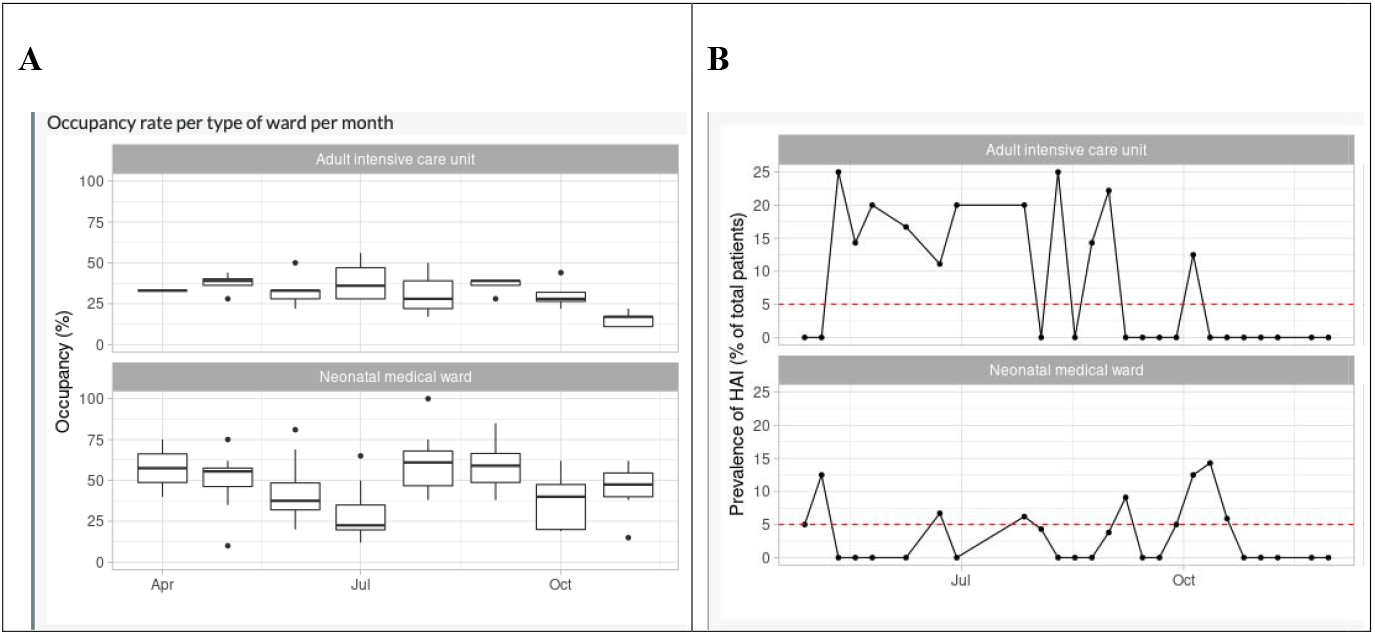
Monthly ward occupancy (A) and bimonthly HAI prevalence (B) in the adult intensive care unit and (for comparison), the neonatal intensive care unit between April and November 2022, illustrating the high and sustained prevalence of HAI in the adult ICU from May. The boxplots represent the distribution of bed occupancy measurements within each month for each ward type. Occupancy was recorded repeatedly within each month; the boxes indicate th median and interquartile range, with whiskers showing the range and points representing outliers. This figure was obtained from the institution’s ACORN dashboard.

Environmental isolates submitted by the hospital’s Infection Prevention and Control team, which were called in to investigate a potential outbreak, included multiple members of the *Enterobacter cloacae* complex (n = 2), *Klebsiella pneumoniae* (n = 7), *Acinetobacter baumannii* (n = 6), and *Pseudomonas putida* (n = 3). Single isolates (n=1) were identified as *Acinetobacter lwoffii, Providencia stuartii, Pseudomonas aeruginosa, Pseudomonas oryzihabitans, Escherichia coli* and *Staphylococcus cohnii*.

Identities of *Acinetobacter* and Enterobacterales were confirmed through whole-genome sequencing. The two *Acinetobacter baumanii* isolates belonged to different STs (ST2833 and ST919) and differed by 52,166 SNPs, ruling out an *Acinetobacter* outbreak. However, the *E. hormaechei* patient isolate, which carried 24 resistance genes, conferring resistance to nine antimicrobial classes, was within 0 to 2 SNPs of two environmental *Enterobacter* isolates, shared resistance patterns with them and had a resistance pattern highly similar to a *Klebsiella pneumoniae* environmental isolate. Following long-read sequencing and hybrid assembly of the *Enterobacter* strains, we additionally reanalysed *Enterobacter* genomes from the same facility [20], including a clonal bloodstream cluster from 2020 which belonged to *E. hormaechei* ST109.

### Whole genome sequence-inferred strain identities, interrelationships and antimicrobial resistance gene (ARG) content

The two confirmed clinical strains of *A. baumannii* were not clonal and did not harbour any detectable plasmid replicons. The three *Enterobacter hormaechei* clinical (UCH_778) and environmental (UCH_710 and UCH_770) isolates, all belonging to ST114, had identical resistance genes and plasmids detected. Six environmental *K. pneumoniae* isolates belonged to five STs. All five *A. baumannii* and *E. hormaechei* isolates were resistant to the carbapenems ertapenem and meropenem. Carbapenem resistance was attributable to *bla*_NDM-1_ genes in both *A. baumannii* isolates and to *bla*_NDM-5_ in the *E. hormaechei* isolates.

Across the dataset, multiple plasmid replicon families were identified, including Col-type plasmids (e.g., Col440I_1, Col440II_1, Col3M_1), IncF variants (IncFIA, IncFIB, IncFIC, IncFII), as well as IncHI1B_1_pNDM-MAR, IncHI2A_1, IncHI2_1, IncR_1, and RepA_1_pKPC-CAV1321. Some isolates carried only one or two replicons, such as UCH-619 (*K. pneumoniae* ST3113), which harbored Col440II_1 and Col440I_1, and UCH-B0249_S1 (*Proteus mirabilis*), which carried a single Col3M replicon. In contrast, isolates such as UCH-935D (*K. pneumoniae* ST395), UCH-943-ECO_S15 (*K. pneumoniae* ST967), and UCH-B0265-KLEB_S10 (*K. pneumoniae* ST147) carried between five and six replicon types, including multiple IncF plasmid families. Several of these strains also carried RepA_1_pKPC-CAV1321, IncFIB(K)_1_Kpn3, or IncFIB(AP001918)_1 (Figure 2). In spite of this heterogenous plasmid landscape, all three *Enterobacter* strains harboured identical plasmid replicons and antimicrobial resistance genes as already mentioned. All three harboured Col440I, IncH12, IncH12A (multi-replicon plasmid commonly observed in IncH plasmids), IncR, and RepA_1_pKPC-CAV1321 replicons, regardless of isolation source. The plasmid replicons and antimicrobial resistance genes identified in the *E. hormaechei* strains were similar to those seen in two out of seven *K. pneumoniae* isolates, suggesting that mobile genetic elements may have been transmitted in the ICU.

**Figure 2:**
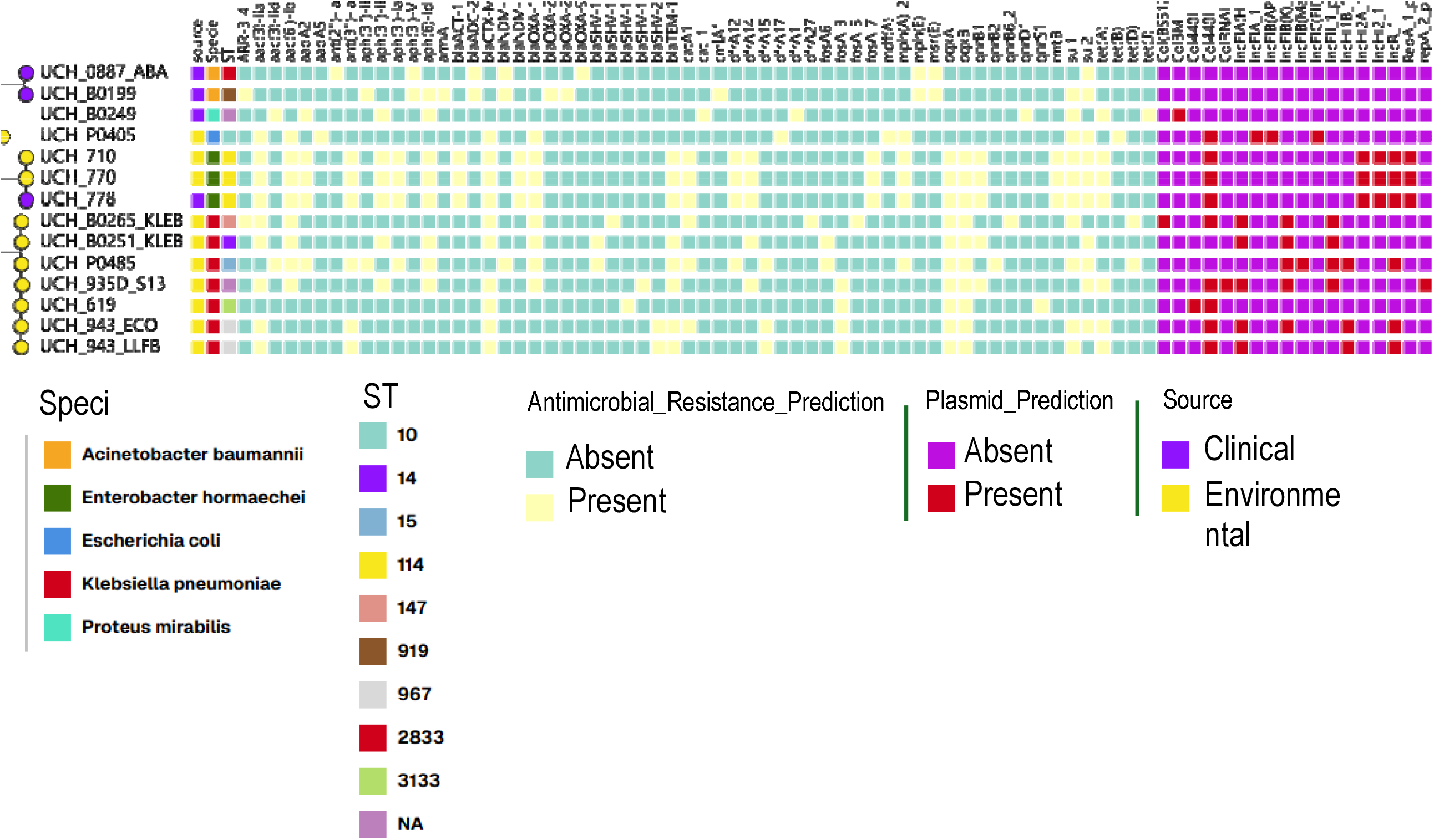
phylogeny of sequenced isolates showing isolate source, plasmid and AMR prediction.

A total of 214 antimicrobial resistance gene hits were identified across all genomes. Of these, 111 (51.9%) exhibited <100% sequence identity and/or coverage relative to reference sequences, indicating the presence of allelic variants or partial gene matches (Supplementary table 2). The number of unique ARGs per isolate ranged between 6 and 24 determinants (Figure 2). Analysis of the 14 isolates revealed the presence of diverse antimicrobial resistance genes across multiple antibiotic classes. The most prevalent genes were *bla*_CTX-M-15_, *oqxA, oqxB, sul1*, and *sul2*, each detected in 10 isolates (71.4%), indicating widespread resistance to β-lactams, sulfonamides, and efflux-mediated drug export. Aminoglycoside resistance genes, including *aac(3)-Iia (*n=9, 64.3%), *aac(6’)-Ib-cr, aph(6)-Id*, and *bla*_OXA-1_ (all n=8, 57.1%), were moderately prevalent. The tetracycline resistance gene *tetA* was detected in 7 isolates (50%). Several additional resistance determinants were identified at lower frequencies (21.4–42.9%), including *aadA2* (n=5, 35.7%), *dfrA14* (n=5, 35.7%), *mphA_2* (n=5, 35.7%), and *qnrB1* (n=5, 35.7%). Carbapenemase genes *bla*_NDM-1_ and *bla*_NDM-5_ were present in 3 isolates each (21.4%). Rare resistance genes (n=1, 7.1%) encompassed *bla*_OXA-_type variants (*bla*_OXA-203_, *bla*_OXA-23_, *bla*_OXA-94_) and additional aminoglycoside, β-lactam, tetracycline, quinolone, macrolide, chloramphenicol, and fosfomycin resistance determinants.

The three *Enterobacter hormaechei* isolates had the highest number of detected resistance genes of all isolates. Genes conferring resistance to at least nine major antimicrobial classes were detected in these isolates: beta-lactams (*bla*_CTX-M_, *bla*_TEM_, *bla*_OXA_), aminoglycosides (*ant, aph*), quinolones (*qnrS1*), tetracyclines (*tet(A), tet(D*)), sulfonamides (*sul1*), trimethoprim (*dfrA12, dfrA1*), macrolides (*mph(A), mph(E), msr(E*)), phenicols (*catA1, cmlA1*), and fosfomycin (*fosA*). The isolates also carried two 16S rRNA methyltransferase genes, *armA* and *rmtB*. Both genes encode enzymes that modify the aminoglycoside binding site on the 16S rRNA, and may confer high-level resistance to all clinically relevant aminoglycosides.

### Genetic context and comparative analysis of *Enterobacter hormaechei* ARGs

Hybrid genome assembly of the three *Enterobacter* strains identified a large plasmid-associated contig carrying multiple antimicrobial resistance determinants and plasmid replicons. A single contig harboured three replicons—IncHI2, IncHI2A, and RepA_pKPC-CAV1321 – yet the plasmid(s) carrying them could not be fully circularized despite hybrid assembly. UCH_770 enabled better plasmid resolution, with the resistance plasmid (253159bp) (Figure 3) separating into two contigs – a major contig containing most resistance genes and replicon content, and a much shorter contig – providing clearer genomic context. Annotation revealed a complex multidrug resistance region encoding resistance to β-lactams (*bla*_CTX-M-15_, *bla*_TEM-1_, *bla*_OXA_), aminoglycosides (*aac(3)-IIe, aac(6*_′_*)-Ib-cr, aph(3*_′_*)-Ib, aph(6)-Id*), sulfonamides (*sul2*), trimethoprim (*dfrA14*), tetracycline (*tet(A*)), chloramphenicol (*catA1, catB3*), and fluoroquinolones (*qnrB1*). These genes were embedded within a mosaic of insertion sequences, Tn3-family transposases, and a class 1 integron (*intI1*), consistent with extensive horizontal gene transfer. This large plasmid in UCH_770 was compared by BLASTn to homologous large plasmids from isolates UCH_710 and UCH_778 and the result revealed extensive sequence similarity across the plasmids, with near-continuous alignment over most of the contig length (Figure 3). However, in UCH_710 and UCH_778, these homologous regions were distributed across multiple contigs rather than a single continuous sequence. The much larger plasmid fragmentation observed in UCH_710 and UCH_778 is attributable to poor long-read sequencing and short average Nanopore read lengths (supplementary table 1).

**Figure 3.**
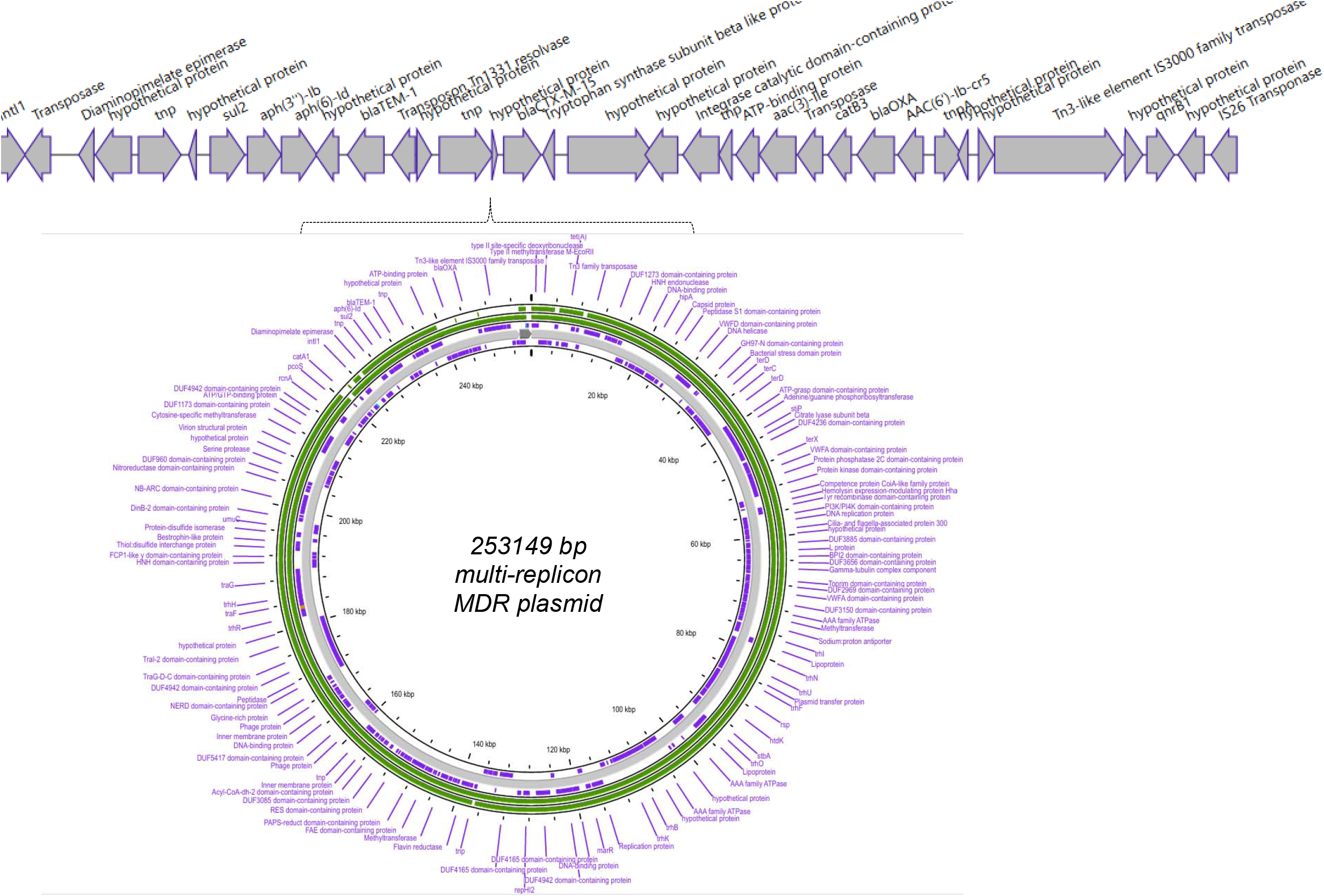
analysis of a large multidrug resistance plasmid contig. The figure illustrates the genetic organization and comparative homology of a large plasmid-ing multiple AMR genes in *Enterobacter hormaechei* strain UCH_770. The top panel shows a linearized gene map of the resistance region, highlighting a The bottom panel presents a circular visualization of the two plasmid contig in UCH_770. The outer annotations (red) indicate the positions of AMR genes. cks represent BLASTn alignments of this contig against homologous large plasmids from isolates UCH_710 and UCH_778. The near-continuous green extensive sequence conservation across these plasmids. Gaps and discontinuities in the BLAST rings correspond to regions that are fragmented across H_710 (outermost green ring) and UCH_778 (innermost green ring), reflecting incomplete plasmid reconstruction due to poor long-read assembly in those

Conversely, IncR plasmids with size between 30557 – 46176 bp, designated pCRE-NDM yielded circularized plasmid (Figure 4). These highly similar plasmids also contained a non-identical but highly conserved region with genes conferring resistance to carbapenems (*bla*_NDM-5_ and *bla*_TEM_), bleomycin (*ble*), sulfonamides (*sul1*), aminoglycosides (*aadA2 and rmbB*), trimethoprim (*dfrA12*), and macrolides (*mph(A), mph(E)*, and *msr(E*)). Further evaluation of the genetic context and comparative analysis of pCRE-NDM plasmid genes in the three *Enterobacter hormaechei* isolates (Figure 5) revealed clustering of the resistance genes within a transposon-rich environment, along with high structural synteny interspersed with localized genomic plasticity. The *bla*_NDM-5_ genetic environment was highly conserved across all three isolates, sharing nearly 100% sequence identity in the core resistance backbone. The environmental isolate (UCH_770) harbored a truncated *bla*_TEM_ gene; however, this had no apparent effect on the resistance phenotype due to the presence of additional β-lactam resistance genes. Most of the differences between the plasmids were due to small hypothetical open reading frames interspersing resistance genes in the resistance cluster, which was replete with insertion sequences, phage sequences, and transposons.

**Figure 4.**
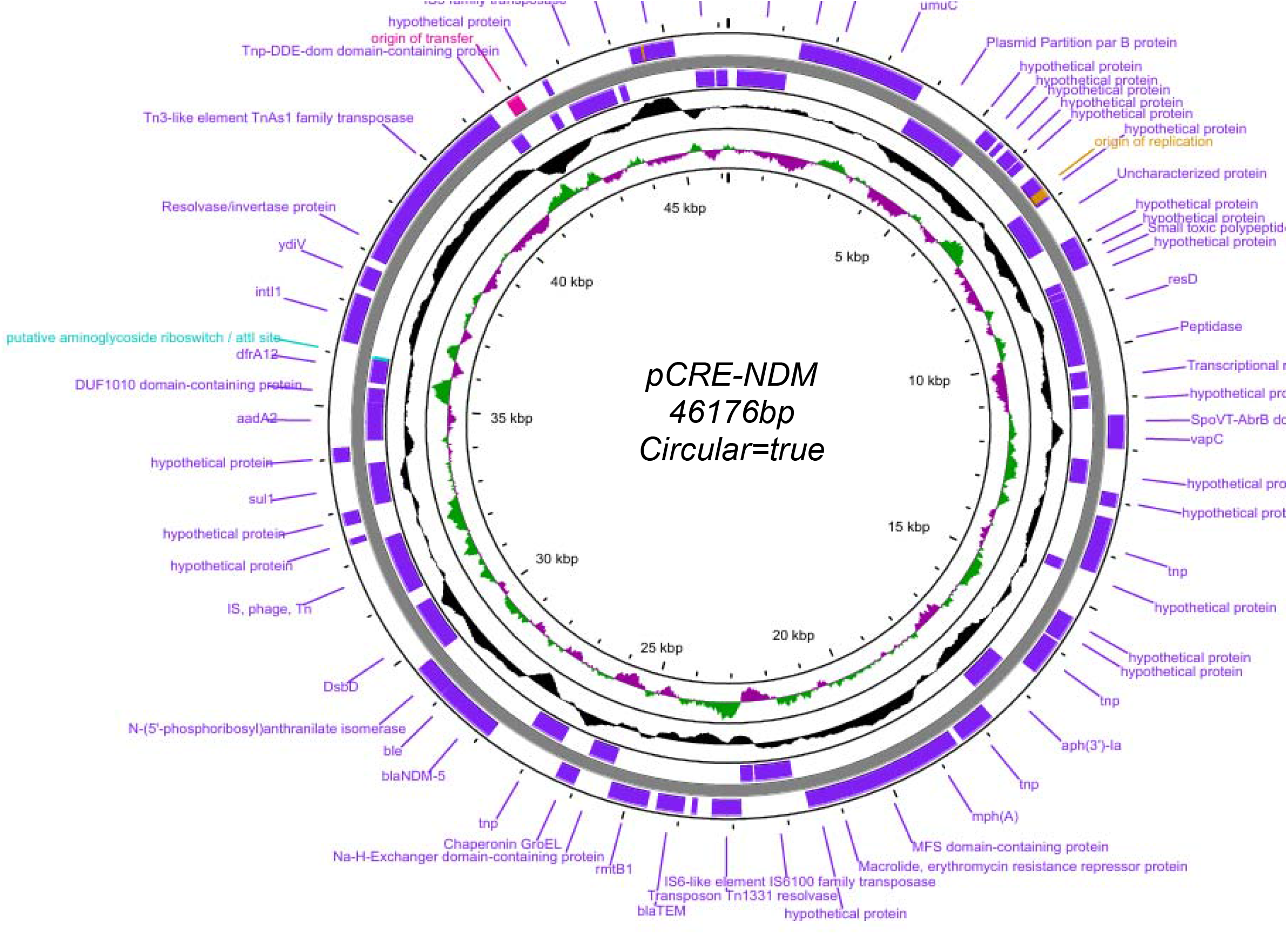
Complete plasmids identified in three clonal *Enterobacter hormaechei* strains. The figure highlights the fully circularized plasmid obtained from the *Enterobacter* strain UCH_770, which represents the complete plasmid structure within this clonal group.

**Figure 5.**
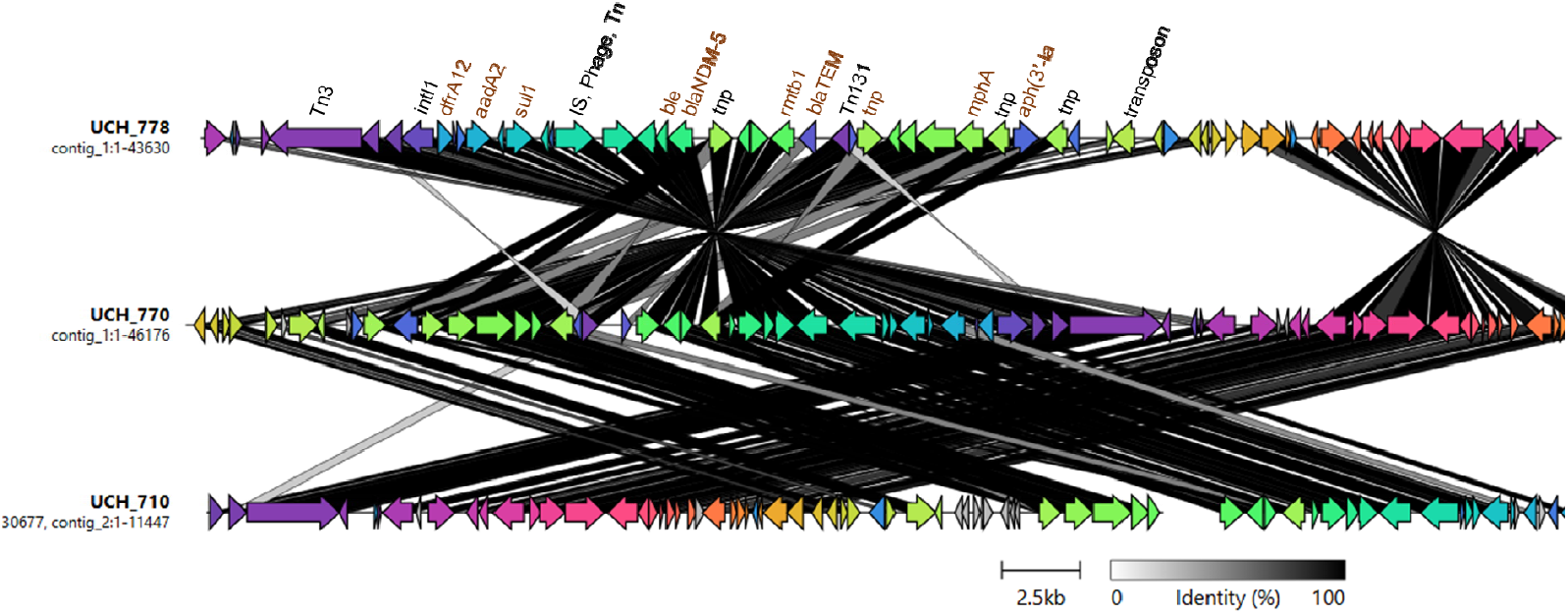
Genetic context and comparative analysis of pCRE-NDM plasmid genes in the three *Enterobacter hormaechei* isolates.

Interestingly, the *K. pneumoniae* ST15 strain (UCH_P0485) isolated from the same ICU ward environment harboured the same resistance genes and plasmid replicons seen in all three *E. hormaechei* strains. Plasmid reconstruction using *E. hormachei* hybrid genome [please specify which strain] as scaffold showed a near identical 46,176 bp plasmid from the *K. pneumoniae* short reads (Supplementary figure 1). Based on this finding, we screened retrospective *Enterobacter* genomes (20) and identified four clonal bloodstream isolates from 2020 belonging to *E. hormaechei* ST109 that carried near identical resistance clusters. The ST109 bloodstream isolates each carried 13 antimicrobial resistance genes (ARGs) and two plasmid replicons. Complete plasmid reconstruction revealed a single 318 kbp circular conjugative plasmid carrying both IncFIB and IncHI1B replicon types. The plasmids had a conserved ∼45 kbp MDR island habouring a cassette carrying *bla*_NDM-5_ alongside other resistance genes such as *bleMBL, mph(A), sul1, dfrA12, aadA2*, and *aph(3’)-Ia*, organized within class 1 integrons and flanked by different insertions sequences and transposase genes. This MDR island was highly similar to the one seen in the ST114 (Figure 6).

**Figure 6.**
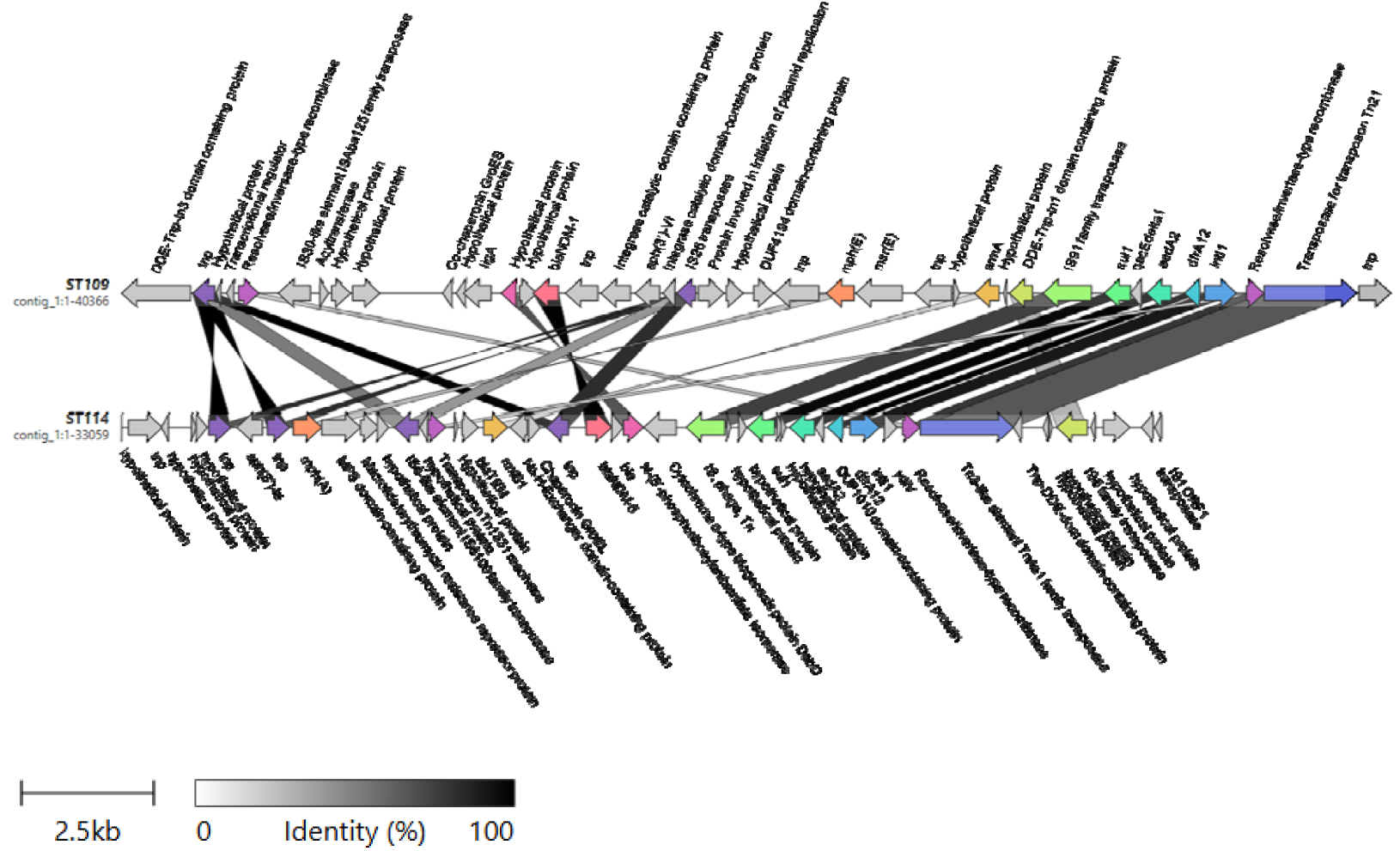
Comparative alignment of the conserved multidrug-resistance (MDR) island found in *E. hormaechei* ST114 and *E. hormaechei* ST109

## DISCUSSION

Healthcare-associated infections are common worldwide and are frequently caused by multidrug-resistant organisms. Although reported less frequently in African countries (21), HAIs are thought to represent a substantial burden, where they may be missed altogether, or, if suspected, their etiologies may not be determined due to inadequate clinical microbiology resources. Even when etiologic agents can be identified, resources for subtyping – necessary to rule in or out outbreaks - may not be available. In this context, we report the results of investigating a suspected outbreak raised by an astute health care worker, in which regular HAI ward surveillance through the ACORN pilot clearly showed a sustained increase in HAIs in the hospital’s intensive care unit. Available clinical microbiology data led to a suspicion of an *Acinetobacter* outbreak because two *Acinetobacter* spp had been cultured within the HAI surge period.

As Nigeria’s AMR surveillance system has centralized whole genome sequencing capacity (22), the hospital was able to request reference laboratory assistance to confirm or rule out an outbreak. In UCH, as in most other Nigerian hospitals, microbiology culture is paid out of pocket by resource-limited patients, resulting in low culture rates. For this reason, recent blood isolates that did not belong to the *Acinetobacter* genus were requested along with isolates suspected to be outbreak strains, as well as all carbapenem-resistant environmental isolates. The broader sampling plan enabled genomic analyses beyond the initially suspected *Acinetobacter* outbreak isolates. This approach enabled generation of new hypotheses centered on mobile genetic elements in the course of ruling out *Acinetobacter* as an outbreak cause. A carbapenemase-gene-carrying mobile element was subsequently identified, following long-read sequencing and hybrid assemblies, in multiple environmental and one clinical isolate.

Of immediate concern to the facility was that the upsurge in infections was associated with ESBL and carbapenemase-producing strains, which were additionally recovered from the ward environment. Studies have documented widespread environmental colonization by ESBL and carbapenemase-producing Enterobacteriaceae and *A. baumannii*, with genetic analyses confirming clonal dissemination within wards and between patients and the environment (23– 25). In this study, the two *Acinetobacter* isolates were from previously reported ST919 and ST2833, neither of which is commonly reported as causes of HAIs in humans. By contrast, the clinical and environmental *Enterobacter* strains isolated belong to frequently reported STs from invasive hospital infections including *E. hormaechei* ST114 (and the previously isolated ST109) as well as notable MDR or hypervirulent high risk clones of *K. pneumoniae*: ST14, ST15, ST147, ST395 and ST967 (26). These strains carried *bla*_NDM-1_, *bla*_NDM-5_, and *bla*_OXA-232_ carbapenemases genes and therefore represented a significant risk to ICU patients, irrespective of whether they were directly involved in the outbreak. In addition to the carbapenemase content, these strains exhibited a complex resistance profile characterized by the presence of multiple beta-lactamase genes. Specifically, the isolates carried several *bla*_SHV_ variants, the broad-spectrum penicillinase *bla*_TEM-1B_, and the extended-spectrum beta-lactamase (ESBL) *bla*_CTX-M-15_. Furthermore, the presence of *bla*_OXA-1_ which is often associated with resistance to beta-lactam/beta-lactamase inhibitor combinations further contributes to a robust MDR phenotype alongside the primary carbapenemases. The mobilization of *bla*_CTX-M_ genes is often facilitated by mobile elements, such as ISCR1 or IS26 structures (27).

The number and conservation of resistance genes detected in the *Enterobacter* isolates were of particular concern. We identified two key contigs in the *Enterobacter* strain (UCH_770). One contig carried three plasmid replicons, IncHI2, IncHI2A, and RepA_pKPC-CAV1321, but despite performing hybrid assembly, this plasmid could not be fully circularized. The second contig corresponded to an IncR plasmid, which was successfully circularized following hybrid assembly. The IncHI2 replicon and partial plasmid sequence, and the IncR plasmid were found in all three *Enterobacter* strains and the *Klebsiella pneumonia* strain. IncH plasmids are well-recognized for their ability to carry multiple AMR genes and to disseminate carbapenem resistance due to their large size and high conjugation potential (28). IncH (especially IncHI2) plasmids are large, stable, but can initially impose a moderate fitness cost that is often ameliorated over time (29). IncR plasmids, although typically non-conjugative (30), are highly stable and frequently occur as multi-replicon plasmids or hybrid plasmids that expand their host range and resistance spectrum and serve as reservoirs for carbapenemase genes (31).

In this study, the *bla*_NDM_ gene and other carbapenemase determinants were located on both IncH and IncR plasmid types. We observed co-localization of multiple ARG with IncHI2/IncHI2A and RepA_pKPC-CAV1321 replicons with different mobile elements on the same contig. The ARG profile demonstrated extensive co-resistance encompassing betalactams, fluoroquinolones, aminoglycosides, sulfonamides, tetracyclines, and phenicols This suggests that exposure to antibiotics outside the carbapenem class could co-select for this plasmid, thereby indirectly maintaining carbapenem resistance genes (32). The *bla*_NDM-5_ gene detected in this study confers resistance to costly (in our setting) last-resort carbapenem antibiotics. The circularized IncR plasmid carried *bla*_NDM-5_ alongside other resistance genes such as *bleMBL, mph(A), sul1, dfrA12, aadA2*, and *aph(3’)-Ia*, frequently organized within class 1 integrons and flanked by mobile elements (e.g., IS26, IS3000, Tn3-like transposons) and this is consistent with reports elsewhere [32] (Figure 4). The truncation of *bla*_TEM-1_ marks a distinct evolutionary divergence between the clinical and environmental *Enterobacter* strains and suggests that the environmental strains have undergone localized genomic remodeling, likely driven by insertion sequence-mediated recombination, which favors the stabilization of high-priority resistance markers like *bla*_NDM-5_ at the expense of redundant or accessory genes.

The genetic architecture, including the size and the presence of *bla*_NDM-5_ in these *Enterobacter hormaechei* strains, is highly suggestive of the globally successful IncH and IncR plasmid family. Fusion events between IncH (IncH12 and IncH12A) plasmids and RepA_pKPC replicons observed in this study have been increasingly reported in recent years, largely driven by mobile genetic elements (33,34). These hybrid plasmids can simultaneously harbor multiple carbapenemase genes and other AMR determinants, enhancing their epidemiological impact (35,36). The co-occurrence of IncH and IncR suggests a plasmid architecture that facilitates the accumulation and mobilization of *bla*_NDM_ and other carbapenem-resistance genes, supporting their epidemiological role in the spread of carbapenem resistance in Enterobacterales. The identical plasmid profiles in both clinical and environmental *E. hormaechei* isolates, and similarity with a plasmid in *Klebsiella* point to a mobile reservoir or resistance genes in the ICU.

The observation of the same resistance island in clinical and environmental *E. hormaechei* ST114 isolates and in a ward isolate of *Klebsiella pneumoniae* ST15, a globally disseminated high-risk clone, strongly suggests horizontal gene transfer as a major driver of its dissemination. The presence of this resistance island within a substantially larger plasmid in earlier-recovered *E. hormaechei* ST109 further supports the hypothesis that the MDR region is a modular and mobile unit capable of reshuffling between plasmids of varying sizes. Such modularity is increasingly recognized as an important mechanism underlying the rapid spread of carbapenem resistance among Enterobacterales (37,38).

Our findings suggest that the use of any major antibiotic class maintains the plasmids and resistance clusters they bear, promoting co-selection and persistence even in the absence of direct carbapenem use. The conjugative potential of these plasmids further enables rapid spread across Enterobacterales, including *Escherichia coli* as well as *K. pneumoniae*, and *Enterobacter* spp as observed in this study. Our observation that plasmids play a central role in mediating carbapenem resistance in *E. hormaechei* aligns with recent findings in other Enterobacterales, emphasizing the contribution of plasmid-mediated transmission to the persistence and spread of carbapenem-resistant strains. For instance, Liu et al. (39) used machine learning to evaluate risk factors for the dissemination of carbapenem-resistant *K. pneumoniae* in neonatal ICU and demonstrated that plasmids were critical for long-term persistence, while short-term transmission was largely driven by patient interactions within healthcare groups.

## Conclusions

This study provides valuable insights into the importance of careful interpretation of suspected outbreak cases in hospital settings. What was initially suspected to be an *Acinetobacter* outbreak was ruled out, but genomic investigation uncovered a carbapenem-resistant *Enterobacter hormaechei* clone isolated from both patients and the hospital environment. A plasmid found within this clone was also detected in a *Klebsiella* environmental isolate. This plasmid bore a cluster that has previously been seen in a resistant *E. hormachei* lineage at the same hospital. This russian doll-like containment for the *bla*_NDM5_ gene identified in this study allows for complex dynamics and dissemination and multiple scales that could be difficult to connect. Carbapenem resistance in hospital Enterobacterales in our setting is driven not only by clonal expansion but by the horizontal dissemination of a highly stable *bla*_NDM_-associated MDR island capable of integrating into diverse plasmid backbones. Our findings underscore the need for careful monitoring of ICU environments, the need to deploy blood culture-based diagnostics within them and justification for genomic support for outbreak investigations. Furthermore, our findings highlight the interconnectedness of clinical and environmental AMR reservoirs and strengthen the case for enhanced surveillance strategies that integrate a One Health perspective to mitigate the spread of resistance across ecological boundaries. It also highlights the importance of genomic surveillance strategies that track mobile resistance elements, and not just bacterial lineages, to inform evidence-based infection prevention and control in low-resource settings.

## Abbreviations

AMR: antimicrobial resistance
ARGs: antimicrobial resistance genes
STs: sequence types
WGS: whole-genome sequencing
MGEs: mobile genetic elements
ICU: intensive care unit
MDR: multidrug resistance
HAI: Hospital acquired infection
CRE: carbapenem-resistant Enterobacterales
CRAB: carbapenem-resistant *Acinetobacter baum*a*nii*
LMICs: low- and middle-income countries
HGT: horizontal gene transfer
ACORN2: A Clinically Oriented Antimicrobial Resistance Network
(API-20E): Analytic Profile Index 20E
CLSI: Clinical and Laboratory Standards Institute
SNP: single nucleotide polymorphism
ESBL: extended-spectrum beta-lactamase

## Acknowledgement

We gratefully acknowledge the clinical and diagnostic laboratory staff at University College Hospital for their contributions to this study.

## Ethical statement

Ethical approval for sample collection, processing and analysis during the 2022 outbreak through ACORN2 surveillance was obtained from the UI/UCH Ethics committee (Approval number UI/EC/21/0754). Approval to use genomes from routine surveillance for research was granted earlier by the same committee (UI/EC/22/0113)

